# UVC-based air disinfection system for rapid inactivation of SARS-CoV-2 present in the air

**DOI:** 10.1101/2022.08.02.502427

**Authors:** Harry Garg, Rajesh P. Ringe, Supankar Das, Suraj Parkash, Bhuwaneshwar Thakur, Rathina Delipan, Ajay Kumar, Kishor Kulkarni, Kanika Bansal, Prabhu B. Patil, Tabish Alam, Nagesh Babu Balam, Chandan Swaroop Meena, Krishan Gopal Thakur, Ashok Kumar, Ashwani Kumar

## Abstract

The novel coronavirus disease 2019 (COVID-19) infections have rapidly spread throughout the world, and the virus has acquired an ability to spread via aerosols even at long distances. Hand washing, face-masking, and social distancing are the primary preventive measures against infections. With mounting scientific evidence, World Health Organisation (WHO) declared COVID-19 an air-borne disease. This ensued the need to disinfect air to reduce the transmission. Ultraviolet C (UVC) comprising the light radiation of 200-280 nm range is a commonly used method for inactivation of pathogens. The heating, ventilation, and air conditioning (HVAC) systems are not beneficial in closed spaces due to poor or no ability to damage circulating viruses. Therefore, standard infection-prevention practices coupled with a strategy to reduce infectious viral load in air substantially might be helpful in reducing virus transmissibility. In this study, we implemented UV light-based strategies to combat COVID-19 and future pandemics. We tested various disinfection protocols by using UVC-based air purification systems and currently installed such a system in workspaces, rushed out places, hospitals and healthcare facilities for surface, air, and water disinfection. In this study, we designed a prototype device to test the dose of UVC required to inactivate SARS-CoV-2 in aerosols and demonstrate that the radiation rapidly destroys the virus in aerosols. The UVC treatment renders the virus non-infectious due to chemical modification of nucleic acid. We also demonstrate that UVC treatment alters the Spike protein conformation that may further affect the infectivity of the virus. We show by using a mathematical model based on the experimental data that UVC-based air disinfection strategy can substantially reduce the risk of virus transmission. The systematic treatment by UVC of air in the closed spaces via ventilation systems could be helpful in reducing the active viral load in the air.

## Introduction

The Severe Acute Respiratory Syndrome Coronavirus-2 (SARS-CoV-2) is responsible for the current pandemic causing millions of infections worldwide and over 2.6 million deaths due to COVID-19 (1). Apart from taking such an enormous toll on human life, the pandemic has imposed unprecedented economic, societal, and healthcare burdens. While mass vaccination drives across the globe are helping to bring back normalcy, there is an urgent need to prevent the virus transmission through aerosols to curb the air-borne transmission to allow the opening of public places and transport such as schools, cinema halls, buses, trains, etc. (2). The attempts to resume the schools, relatively tightly packed and crowded offices, without adequate measures to prevent virus transmission are prime reasons for continued outbreaks (3). Many methods are being explored for the effective disinfection of materials and common-touch surfaces, such as alcohol-based disinfectants, copper surfaces, and soaps. However, there is no effective and safe method that can be deployed in closed workplaces to inactivate the SARS-CoV-2 virus in the air. The face masks are particularly effective in restricting transmission through aerosols and prevent the spread (4). However, the common breach in face-mask practices is causing transmission of the virus within small groups and bigger gatherings. UV radiation is known to effectively kill bacteria and viruses (5) and recent studies have shown that this method is effective in inactivating SARS-CoV-2 virus(6). The UV light causes the formation of pyrimidine dimers in the genetic material and inhibits transcription and replication(7). Therefore, the targeted damage of the essential component of the microbes could be an excellent strategy to attenuate the infectivity of viruses, especially the ones which transmit rapidly, such as SARS-CoV-2.

The WHO declared in May 2021 that SARS-CoV-2 transmission is not only by common touch, close contact but also through air. Indeed, SARS-CoV-2 was detected in our studies in air samples near hospitals even earlier (8). The air-born transmission has serious implications on the control measures we can deploy in place and emphasizes the reduction of viral load in air. In this study, we explore the UV radiation for its ability to inactivate SARS-CoV-2 present in the aerosols. We study the inactivation kinetics by using different dosages of UV to identify the dosage required to achieve maximum virus inactivation. Devices using UV-C light for sanitization of air within air-ducts of HVAC systems and circulating units for use within occupied spaces were designed, developed and validated for delivering the relevant viricidal doses of UV-C light to the air and maintaining the air change rates required by the ventilation guidelines. Further, a risk assessment model to quantize the risk reduction was developed and implemented where it was found that the use of UV –C light irradiation to sanitize air can result in the reduction of the risk of infection in the occupied spaces by up to 90%.

## Results

### Design of ultraviolet C (UVC) Disinfection system

Based on the market requirements, two types of systems were used – i) in-duct UVC disinfection system and ii) standalone air circulating UV-C disinfection system. These designs are described below.

### In-duct UVC disinfection system

In-duct UV systems work for the purpose of inactivating microbes in airstream in a building or zonal ventilation system. The UV lamp is key component and the design and optimization of in-duct UV systems revolves around various features of it. The in-duct system also depends on the output characterization, required UV dose for microbicidal activity, and energy consumption evaluations of UV systems. The in-duct system we designed consisted of UV lamps, fixtures, and ballasts. In the chamber, the access to the air by UV radiation is uniform and so the UV lamps could be fixed at any location including air handling unit (AHU) (**Figure 1**). While designing the systems, we placed the lamp fixtures and ballasts either internally or external to the ductwork. The drop in pressure with external fitting was relatively less than internal fitting. In any case, the drop in pressure associated with UV radiation was only marginal when the velocity of air was within the normal limits of 2–3 m/s (400–600 fpm). By applying these parameters, the modular systems were made and installed in ductwork. The UV radiation may not disinfect the air in one encounter but recirculation of air from the room via the system will increase such encounters and give multiple UV doses to airborne microorganisms for maximum disinfection. Therefore, the re-circulation systems are more efficient than single-pass systems in which air is not recirculated. The characteristics of an air stream that can impact the design are relative humidity (RH), temperature, and air velocity. However, air temperature has a negligible impact on microbial susceptibility in general although membrane viruses such as coronavirus are more susceptible to higher temperature because of rapid drying of aerosols in the hotter environments. The RH factor is compensated as the UV systems are installed after the conditioned air and before the delivery in user place.

**Figure 1:**
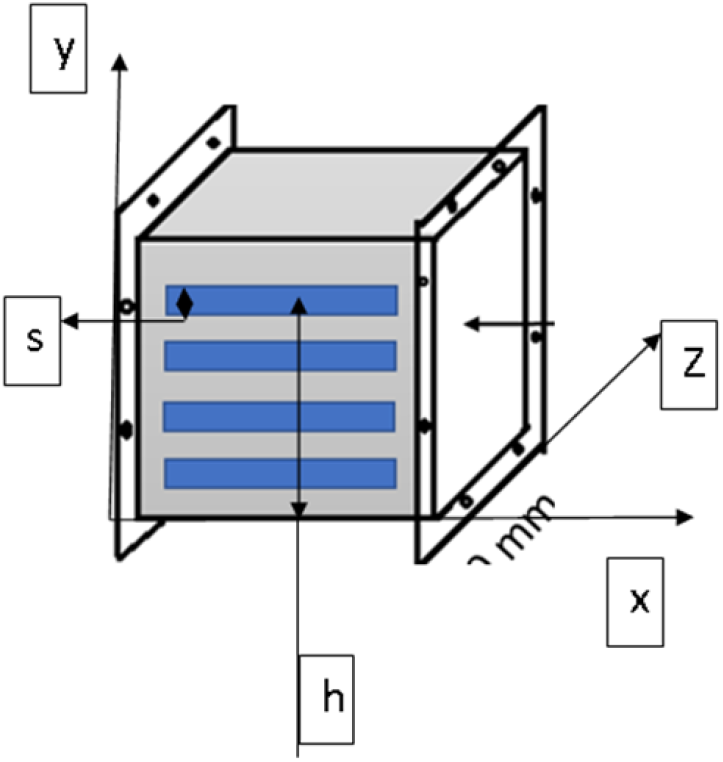
Configuration of induct UV-C disinfection system. Induct UV-C air disinfection system in a cartesian coordinate system with flow in x direction, Illumination in z-direction and source installation in y-direction. The scale is in mm.

In addition to this, duct disinfection system depends upon following parameters i) duct dimensions ii) duct materials iii) air temperature iv) amount of recirculated air v) amount of fresh air in recirculation, and vi) flow rate. The different parameters of the proposed systems were X: Length of the source in X-axis, Y: height of the source surface in Y-axis, Z: target surface in Z-axis, and h: total installation height. The UV intensity was evaluated from these parameters considering the flow rate. The required parameters such as flow rate, exposure time, no of air changes, installation space, and UV intensity were optimized to get the required dosages for deactivation of the virus and maintain the constant fluence rate in the duct. The design of the induct UV-C system (unidirectional or bi-directional) is shown in Figure 2. It shows an enclosed space with source to target illumination surface in z direction and the system was used in the closed indoor space.

**Figure 2:**
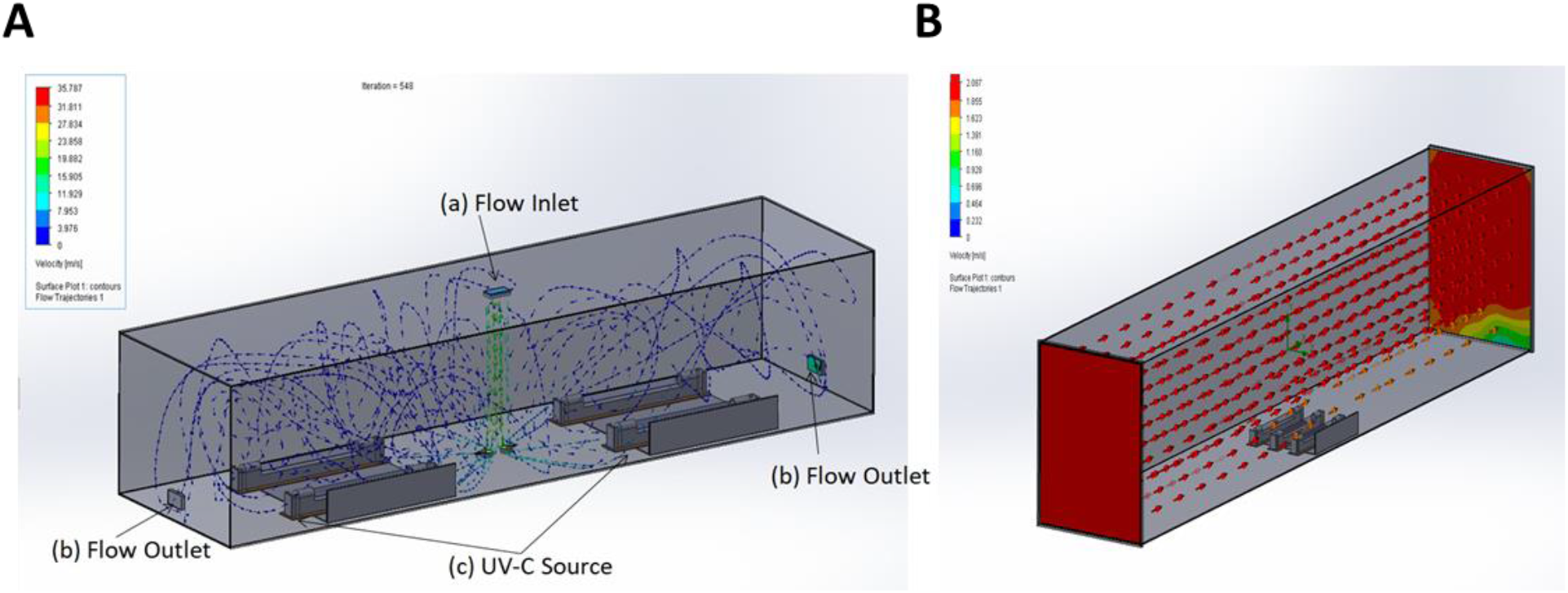
Large size duct with UV-C air disinfection system A) bi-directional flow system with (a) flow inlet at top as infected/recirculated air enters, treated in the duct within volumetric space and disinfected delivery of air exhaust (b) left and right side and (c) UV source fixtures. The blue dotted lines show the flow trajectory of air. B) unidirectional flow system with the UVC source at bottom and air being treated in disinfection volume using. The scale shows the velocity in m/s

The principal design objective for an in-duct UV-C air disinfection system is to create UV energy distribution uniformly throughout a specified length of the duct or air-handling unit (AHU) to deliver the appropriate UV dose to bacteria/virus/aerosol particle in the air moving through the irradiated zone with minimum system power as shown in figure 3. Enhancing the overall reflectivity of the inside of the air handler or air duct improved UV-C system performance by reflecting UV-C energy back into the irradiated zone, thus increasing the effective UV dose and maintain the constant fluency rate. The flow rate was optimized and the change in air flow rate due to the obstruction of the UV source in the flow was evaluated, which was negligible in the design. The designed and fabricated duct is shown in figure 4.

**Figure 3:**
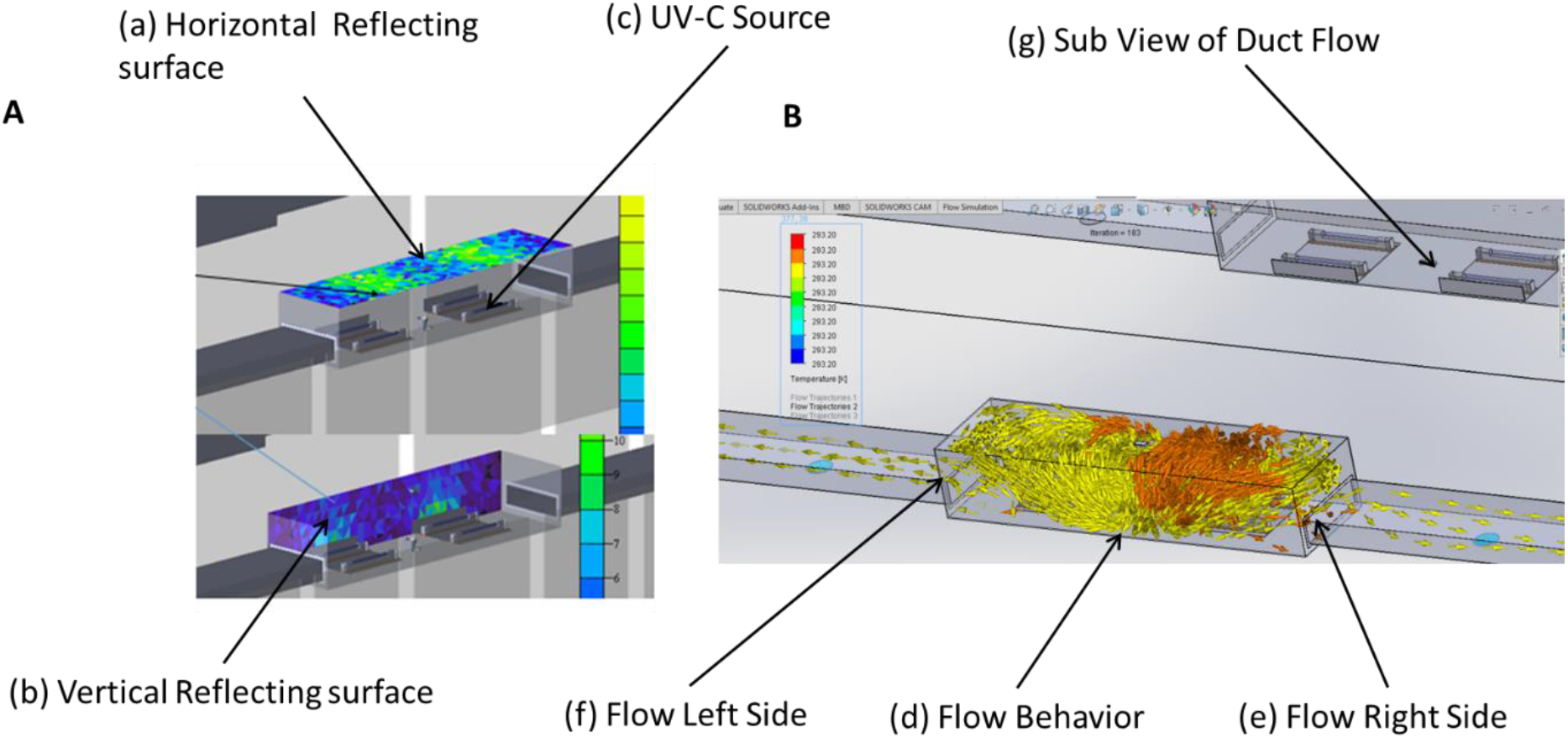
Disinfection of air in flow ducts considering irradiance plot and flow behavior A) The irradiance plot of duct on target surface in horizontal (a), vertical surface (b), and UVC source (c). B) bidirectional flow behavior of air from the center of duct.

**Figure 4:**
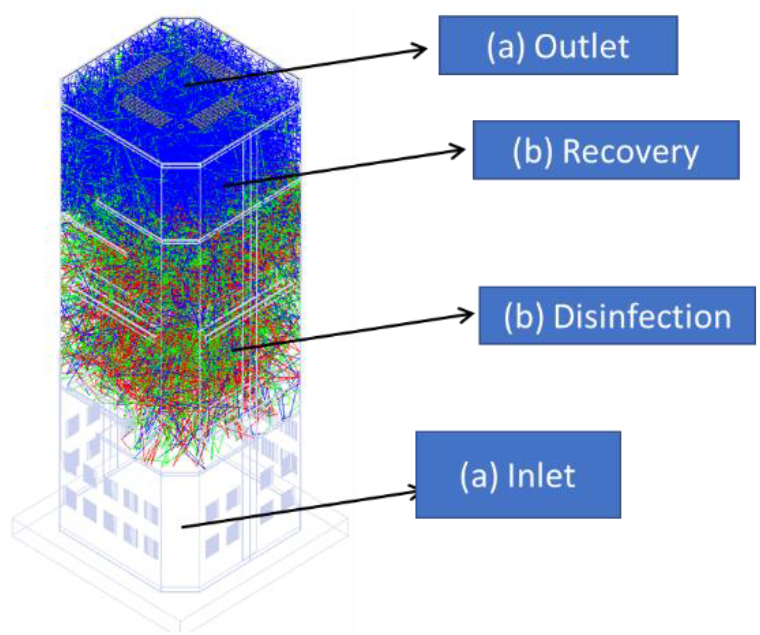
Standalone Air disinfection system with flow in vertical direction. Shown is the inlet for air (a), UVC treatment compartment (b), the recovery area (c), and outlet for the disinfected air (d).

### 1.1 Standalone recirculation Air disinfection

The recirculation UV disinfection systems were installed in the smaller workplaces, each consisted of UV lamps and fixtures in a housing containing a blower. The airflow in recirculation units were in the range of 1.4–14 m^3^/min (50–500 cfm) and were suitable for small rooms or apartments only. Many recirculation units were portable and could be positioned on the floor, tables, mounted on wall or ceiling. Room recirculation units and upper air systems were installed to augment in-duct systems or where in-duct installation was not feasible. The prototype stand-alone unit was tested under laboratory condition and based on the results these units are currently installed for application in the real-world scenario. Due to requirements of compact size, the internal volume of recirculation units do not usually allow extended exposure times to deliver the viricidal dosages during the transit of air through them. Hence, the units were carefully designed to create enough serpentine paths through the circulating devices within the volume irradiated by the UV –C light so as to deliver the viricidal dosages while the units remained compact and portable. The air that goes into the unit comes out sanitized to the extent of over 99% of the virus load reduction.

Both the in-duct and the standalone systems were designed for applications ranging from very small volume and low flow rate to high volumes and high flow rates for different applications such as lifts, toilets, classrooms, workspaces, offices, meeting halls and auditorium etc. The details of the different systems are as shown in table 1.

**Table 1:** Types of UV-C disinfection systems

| S. No | Parameters | In-duct UV-C disinfection systems for HVAC buildings | In-duct UV-C disinfection systems for HVAC buses | Standalone UV-C disinfection system |
| --- | --- | --- | --- | --- |
| i. | Flow rate (CFM) | >30,000 | <3000 | <300 |
| ii. | Applicable space(ft<sup>2</sup>) | >1000 | <350(customizable) | <150 |
| iii. | Shape and size | Rectangular<br>>300mm | Rectangular<br><300mm | Rectangular<br><1500mm |
| iv. | Wattage(W) | >100 | <100 | <70 |
| v. | Weight | <2.5 Kg | <1.2 Kg | <3.0 Kg |
| vi. | UV source(nm) | 254 | 254 | 254 |
| vii. | Length of the delivery duct(m) | >10 | <2 | <0.5 |
| viii. | Exposure time (sec) | 1.1 | 1.14 | 1.1 |
| ix. | Dosages | >1.6mJ/cm<sup>2</sup> | >1.3mJ/cm<sup>2</sup> | >1.3mJ/cm<sup>2</sup> |

### Design of a standalone system to measure the effect of UVC on SAS-CoV-2 survival in air

A hermetically sealed chamber was designed to study the dosage of UVC required to inactivate SARS-CoV-2 present in aerosols. The volume of the aerosol generation and the volume of the UV treatment are interrelated. The aerosols were created from virus suspension by using the nebulizer which produces mist by creating low-pressure zone at the surface of the liquid which pulls up fine droplets from the liquid surface. The shape of the chamber was designed to be a rectangular with the dimension of 56×41×31 cm to ensure the uniform distribution of aerosols (Figure 5B and 6B). The inlet and outlet of the chamber were customized to fit to the nebulizer pipe and sample collection tube respectively. The regulator was installed to control the UV-C intensity. The air filtration unit was connected to the chamber to suction the air from the chamber and a filter was placed in the outlet pipe to collect the aerosols on the surface of the filter membrane.

**Figure 5:**
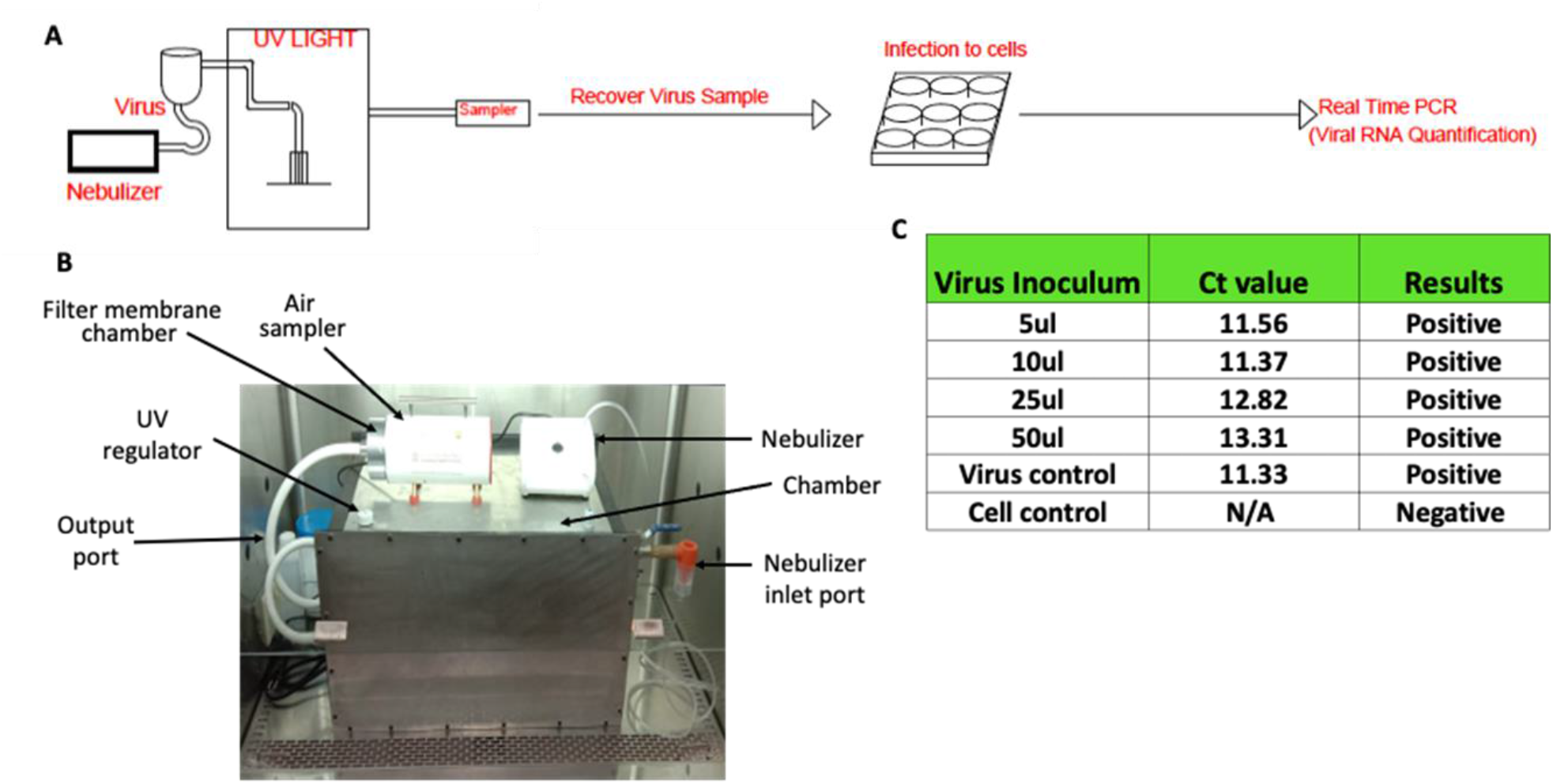
Workflow of SARS-CoV-2 aerosol generation and detection of virus in air. A) The schematic of aerosol generation, virus sampling, infection and detection of virus growth B) The UVC treatment and air sampling device C) The effect of gelatin on the viability of SARS-CoV-2. The virus supernatant was applied onto gelatin membrane and recovered by dissolving membrane in PBS to assess infectivity in Vero-TMPRSS2 cells. The Ct values for original virus supernatant (virus control) and gelatin-recovered virus are indicated in the table.

**Figure 6:**
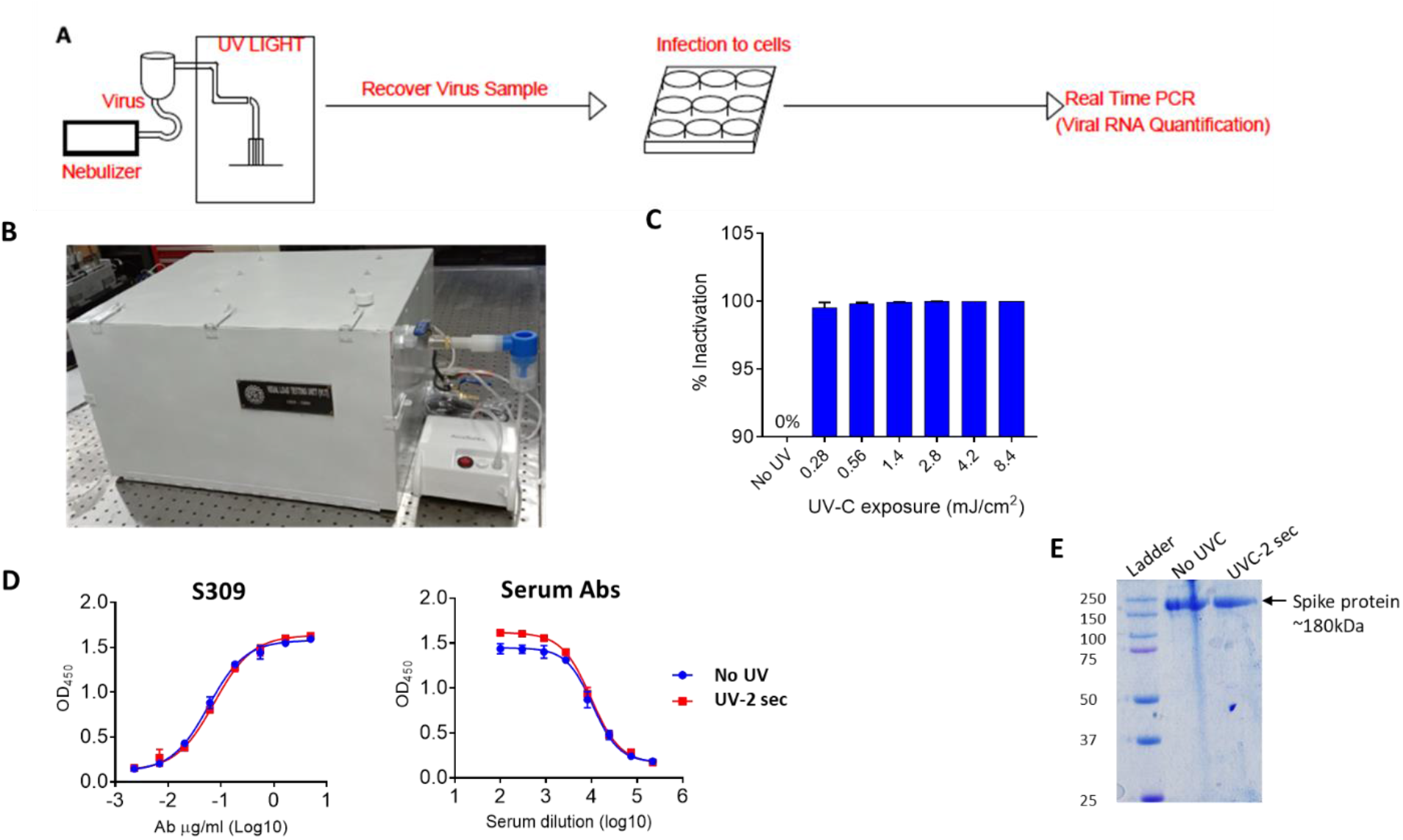
Inactivation of SARS-CoV-2 present in air: A) The schematic of aerosol generation virus sampling, infection and detection of virus growth B) The stand-alone UVC-based device without the air sampling connector to sample air containing aerosolized virus inside the chamber. C) Inactivation of SARS-CoV-2 by UV treatment. The recovered virus aerosols from quartz tube after UV treatment were resuspended in growth medium and used for infection of Vero-TMPRSS2 cells for 48 hours. Cell supernatants were analysed forviral RNA. Percent inactivation of virus at various UV-C exposures with reference to no UV treatment was recorded. D) The binding of S309 or serum antibodies to spike protein is shown in dose-dependent manner. E) The spike protein with or without UVC treatment in SDS-PAGE.

### Testing the efficiency of Induct UVC air disinfection system against aerosolized SARS-CoV-2

A number of different methods are used to entrap the airborne viruses. These include impactors, electrostatic precipitators, filters, etc. (9). The transmission of COVID-19 has been primarily estimated through collecting specified volumes of air on filters followed by estimation of viral load through quantitative real-time polymerase chain reaction (qRT-PCR) (10, 11). However, such methods only look at the presence of virus particles in the air samples which does not provide any information on infectious dose. Since our primary goal was to evaluate the effect of UV radiation on the infectivity of air borne SARS-CoV-2, we designed an air sampling chamber as described above to entrap air-borne SARS-CoV-2 and assess the infectivity after UV irradiation (design of device and experimental set-up is shown in Figure 5A and actual device is shown in Figure 5B). For each cycle of aerosol generation, virus suspension containing 1.5 × 10^7^ pfu was nebulized into the chamber. Two minutes were sufficient to produce aerosols from approximately 90% of the loaded virus stock in the nebulizer. The nebulized SARS-CoV-2 was either collected directly (control) or exposed to UV light and then collected by trapping them onto a gelatin filter using an air sampling device that channeled the air through a gelatin filter (Figure 5B). The filtration unit was turned on with the speed of 100 L/min of air to collect virus particles on the filter. The gelatin filter was solubilized in DMEM and then used to infect Vero-E6-TMPRSS2 cells in a 48-well plate, each well containing 4 × 10^4^ cells. The Vero-E6-TMPRSS2 expresses a moderate level of ACE-2 but high TMPRSS2, which has been shown to have increased SARS-CoV-2 infectivity by 100-folds(12). The infection by collected sample from the membrane was performed for 75 minutes, after which cells were washed, and grown in fresh growth medium supplemented with 5% FBS for 48-hours. Virus particles were quantified by examining the viral RNA in culture media. After 48 hours of infection, the viral RNA was not detected in control as well as UV-treated samples. To test whether gelatin had any virus inhibitory property, virus suspension (10^6^ pfu/ml stock) was directly added to the gelatin filters and incubated for 10 minutes. After incubation membrane was solubilized in DMEM as described above and used for the infection of cells. The viral genomic RNA was detected in the culture supernatants indicating that virus infection and replication took place. replication we were able to culture infectious SARS-CoV-2 (Figure 5C). Collectively, these findings indicate that SARS-CoV-2 viruses are probably quite fragile and susceptible to air-drying and collision onto the solid surfaces.

### Testing the efficiency of standalone UVC air disinfection system against SARS-CoV-2 virus

As mentioned above, the traditional device such as an air sampler was not suitable for the analysis of infectivity of SARS-CoV-2 due to mechanical disruption of virus particle by collision with the filter. Therefore, we aimed to develop a tool wherein we could expose the SARS-CoV-2 to UV and then collect the virus to analyze its infectivity (Figure 6A). We were specifically interested in using UV irradiation for disinfection of SARS-CoV-2 since it has several advantages over existing chemical and conventional disinfectants, such as negligible emergence of resistance, and it does not chemically affect the material. To circumvent the issues related to the trapping of viral particles through an air filtration device, we took inspiration from the natural mode of SARS-CoV-2 transmission. The droplet nuclei containing SARS-CoV-2 are usually inhaled by healthy individuals and deposited inside the airway system. To create a new device to monitor the infectivity of viral particles, we utilized aerosol’s tendency to settle on the surfaces such as glass and steel to trap the viruses (13). SARS-CoV-2 was nebulized, and then the nebulized particles were released into a quartz tube in an air-tight aerosol chamber connected to the nebulizer (Figure 6B). This device was placed inside a biosafety cabinet to ensure the containment of aerosolized SARS-CoV-2. The device was equipped with a 30 mW UV light tube enabling exposure of the trapped viruses in the center of the device. After UV exposure, the quartz tube was rinsed with cell culture media to recover the viruses. Since we were unable to recover viruses directly by passing aerosolized air through a gelatin filter in the induct UVC disinfection system, here we used quartz tubes to collect the aerosols and then exposed to UV. The open quartz tubes were placed inside the chamber to collect the aerosol sample. The suspension containing 1.5 × 10^7^ pfu virus was aerosolized for two minutes as described above and then exposed to UV light at different doses ranging from 2.8 mJ/cm^2^ to 16.8 mJ/cm^2^. UV dosage of 0.28 mJ/cm^2^ was sufficient to inactivate 99.2% SARS-CoV-2 whereas UV dosage of 0.56 mJ/cm^2^resulted in 99.8% inactivation of SAR-CoV-2 (Figure 6C). Thus, we concluded that 0.28 mJ/cm^2^ is sufficient to inactivate the aerosolized virus meaningfully. The disinfection of virus is most likely due to chemical modification of genomic RNA but we also investigated whether UVC has any effects on Spike protein of SARS-CoV-2. To this end, we used SARS-CoV-2 spike protein ectodomain (spike-6P) stabilized in pre-fusion state. The UVC irradiation equivalent to 0.28 mJ/cm^2^ enhanced the binding of serum antibodies but not the S309 antibody which recognized the conserved epitope in RBD (Figure 6D). The UVC treatment did not break the peptide chain and only one intact band appeared on SDS-PAGE (Figure 6E). This data suggested that UV radiation alters the conformation of spike protein and may thus affect the entry of the virus. In summary, the above-described observations suggest that UV radiation can efficiently inactivate the air-borne SARS-CoV-2 and it could be used for disinfection of air.

### Risk analysis of infection and reduction of risk

The UV-based air disinfection system resulted in rapid loss of infectious SARS-CoV-2 and based on this data we next assessed the reduction of transmission risk when this system is used in real world scenario. Transmission of pathogenic microorganisms is a complicated process. This comprises pathogen features, the number of particles produced in a potentially pathogenic host, how effectively the pathogen survives or remains viable outside that host, and the immune system of a person who is exposed to the pathogen. For decades, the Wells-Riley model has been primarily employed for this purpose (14).

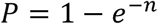

Where, P is the probability of infection for susceptible persons and n is the number of quanta inhaled. The quanta inhaled is influenced by the average quanta concentration (C_avg_, quanta/m^3^), volumetric breathing rate of an occupant (Q_b_, m^3^/h), and the duration of the occupancy (D,h).

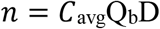

The airborne quanta concentration rises with time from zero to one minus exponential. The time dependent airborne concentration of infection quanta depends on loss rate coefficients which is a sum of ventilation rate, deposition on to surfaces, virus decay and filtration or air disinfection efficiency(15).

To quantify the impact of the virus concentrations, we considered an example of a secondary school class room which consists of 40 students and a faculty. Let us consider faculty is infected with SARS-CoV-2 assuming no susceptible student is wearing mask and are un-vaccinated. The breathing rate of the faculty as per the taking activity is 1.1 (15) and the quanta emission rates for this activity as 9.7 (16). The recommended rate of air circulation for school building is 5 air changes per hours (17). The surface deposition loss rate considered as 0.3 (1/h) (18, 19). Fears et al. (20) observed no virus decay in virus-containing aerosol for 16 hours at 53% relative humidity, but van Doremalen et al. (13) calculated the half-life of airborne SARS-CoV-2 to be 1.1 h, corresponding to a decay rate k= ln(2)/t1/2 of 0.63 1/h.

As per the investigation the efficiency of the UV dosage of 0.28 mJ/cm^2^ leads to a 99.2% while UV dosage exposure of 0.56 mJ/cm^2^ resulted in 99.8% inactivation of SAR-CoV-2. Consider the UV-C disinfection system efficiency as 99.5% as an average of two, the risk of infection and reduction in risk calculation for class room with 6 hours of operation is presented in Figure 9. Without UV disinfection, after 6 hours of occupancy in the classroom, 13 students out of 40 could become infected with SARS-CoV-2 (**Figure 7**). In comparison, after implementing a UV disinfection solution, the risk of infection is reduced by 90%.

**Figure 7:**
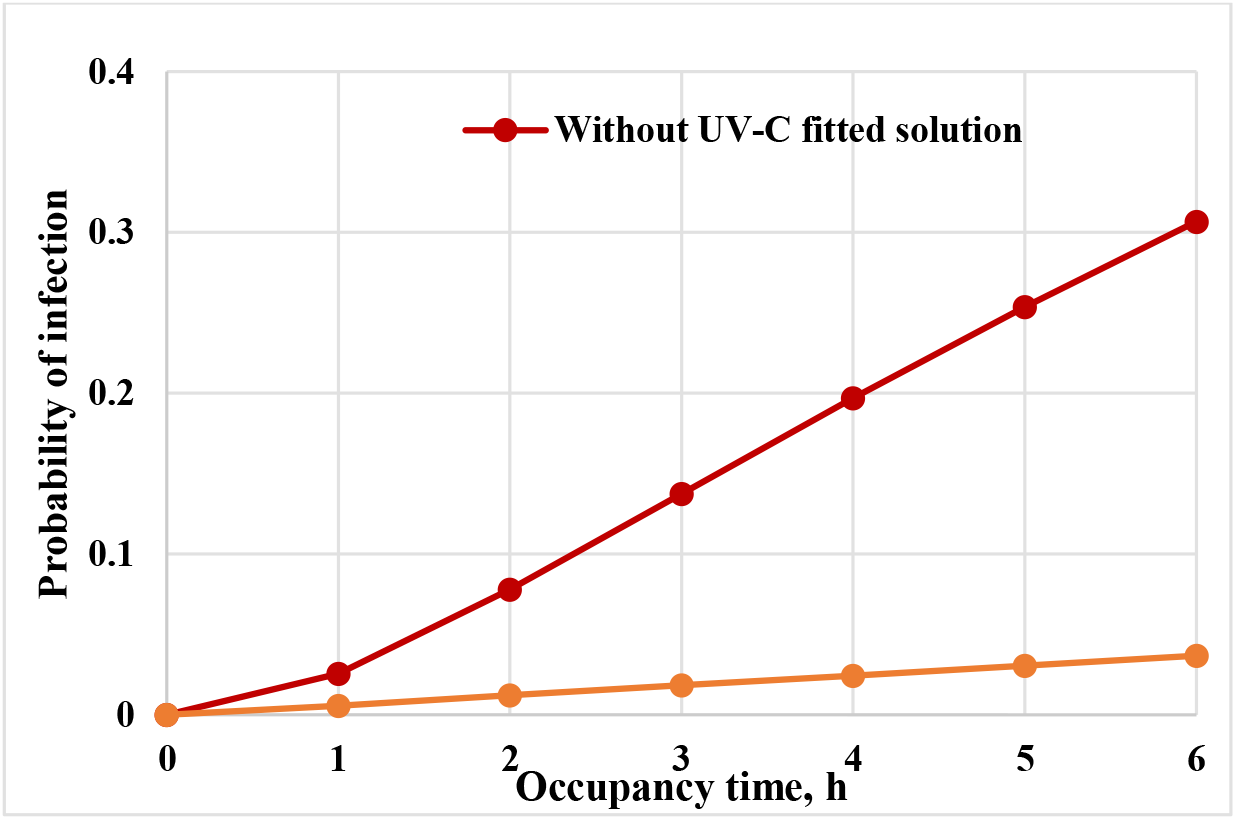
The mathematical model of the risk reduction by UV-C treatment of air. Based on the results of SARS-CoV-2 virus inactivation using stand-alone device the risk reduction in the class room was modelled. Shown is the % reduction of risk with the function of exposure time.

## Materials and Methods

### Design of UV Disinfection system

Both the systems are developed using commercial bill of material (BOM). Certified UVC sources, Aluminum plates with surface treatment, commercial grade fasteners and fittings were used. The UV lamps used in the system were of 254nm wavelength and were obtained from commercial source. The qualified power supplies, Teflon wiring with extra sleeves was used as safety measure. The developed system also met all the commercial safety, environmental and electromagnetic induction (EMI) directives. The systems were robust in design and had satisfactory performance to inactivate coronavirus at the installed places. All the parameters considered for designing the system are listed in table 1.

Both the induct UVC air disinfection system and the standalone air disinfection system are designed according to the required optimum dosages, flow rates and the duct dimensions. The system was designed in such a way that it does not affect the flow rates through the delivery duct and the UVC systems were mounted along the side wall in induct UVC air disinfection system. The Induct system comes in two configurations i.e extraction & retraction type mechanism (refer figure 1 & 2) for the overhead delivery ducts and the fixed type (refer figure 2 and 4) for the separated delivery ducts. The standalone system (figure 5) is used in public places with the human life and has a fixed UVC sources and is designed to meet the standard COVID ventilation guidelines and 10 ACH (air changes per hour). Both the systems were adequately sealed and UV leakage protection was ensured through the optimum sealing. In order to increase the effectiveness of the UVC source, the aluminum polished plates were used and the irradiated zone had high finish aluminum surface for efficient delivery.

### SARS-CoV-2 virus preparation

All the experiments involving the handling of the SARS-CoV-2 virus were performed in the BSL3 facility at CSIR-Institute of Microbial Technology, Chandigarh, according to the institutional biosafety guidelines (IBSC approval no CSIR/IMTECH/IBSC/2020/J23) and institutional ethics guidelines (IEC May 2020#2). The SARS-CoV-2 strain used in the study was isolated from an Indian patient and cultured using the VeroE6 cell line as per the established methods (21, 22). The SARS-CoV-2 was confirmed by whole-genome sequencing and the sequence was submitted to gene bank (Accession # EPI_ISL_11450498). An aliquot of the virus from passage-1 was used to inoculate the 25mm cell culture flask containing Vero E6 cells with 80-90% confluent cells in 5 mL medium. The virus growth was monitored regularly. After extensive cytopathic effect, virus suspension was harvested, clarified and aliquoted in microcentrifuge tubes for long storage at -80°C until further use. The viral load was estimated by quantitative real-time PCR (GCC Biotech) by using 10 µl sample diluted in 100 µl growth medium. Same virus stock was used for all the experiments.

### Aerosol generation, sample collection and infection of cells by aerosol samples

Before setting up the experiments, the device was cleaned and disinfected using 70% ethanol. The working 0.5mL virus stock was freshly thawed for aerosolization. The sterile quartz tube was placed inside the UV chamber and the chamber was closed from all sides. 0.5 mL virus solution was loaded in the sample holding cup of the nebulizer and the mouthpiece was then attached to the inlet tube of the UV device. The aerosols were created and allowed to accumulate in the device chamber for 2 minutes. UV irradiation was done for the required period of time by using a UV light switch outside the bio-safety cabinet. After UV treatment, the quartz tube was removed carefully from the UV chamber, and immediately 0.5mL growth media was added to rinse the surface of the tube. 50 µL sample was then used to infect the cells in a 48-well tissue culture plate containing 5× 10^4^Vero-E6-TMPRSS2 (JCRB1819) cells per well(12). The infection was done for 1 hour with intermittent swirling. After incubation, cells were washed with 200 µL sterile PBS, and a fresh 200 µL growth medium was added to the wells, and the plate was further incubated for 24 hours at 37^0^C in the atmosphere of 5% CO_2_.

### Quantitative real-time PCR (qRT-PCR): The Vero cell supernatants were harvested from the test plate after required incubation

RNA was isolated from 140ul supernatant for qRT-PCR-based analysis. The RNA isolation by Qiagen RNA isolation kit and qRT-PCR by GCC Biotech was performed by using the manufacturer’s instructions. The qRT-PCR was performed by using Bio-Rad system CFX96 Real-Time System. The extent of virus inactivation was calculated by quantifying viral RNA in respective virus cultures from UV-treated aerosols and non-treated aerosols.

### SARS-CoV-2 spike protein expression

expi293 cells were cultured in 30ml Erlenmeyer flasks in freestyle expression media in humidified incubator with 5% CO_2_ at 37°C temperature. The shaking speed was 130rpm/min. The cells were seeded 1 day before transfection to achieve 3×10^6^ cells/ml at the time of transfection. The cells were transfected by using polyethyleneimine (PEI) transfection reagent (Polysciences). Transfected cells were further incubated at constant shaking speed for 4 days before harvesting the culture supernatant. The supernatant was filtered through 0.45u filters and passed slowly through HisPur Ni-NTA resin column (Thermofisher Scientific) with the speed of ∼0.1 ml/minute. The nickel beads were washed twice with washing buffer (NaCl) followed by elution with Imidazole (150mM). The eluate was then dialyzed overnight in PBS and protein was concentrated by using Amicon Pro centrifugal filter (Millipore). The protein was quantified by using BCA Protein Assay Kit (Thermofisher Scientific).

### SDS-PAGE

2ug protein was loaded on 10% SDS-gel followed by staining with Coomassie brilliant blue dye.

### Enzyme-linked Immunosorbent Assay (ELISA)

1ug/ml spike protein in PBS was coated overnight in 96-well plate followed by washing and blocking the plate with PBS containing 5% FBS plus 1% skimmed milk for 1 hour. After blocking, S309 or vaccine serum was added in serial dilution and incubated for 2 hours at room temperature. The antibodies were washed thrice using PBS and anti-human secondary antibody diluted as 1:3000 was added for 1 hour. The plate was washed four times with PBS containing 0.1% tween20 followed by addition of TMB substrate. The reaction was stopped by adding 0.3M H_2_SO_4_. The optical density was recorded by reading the plate at 450nm.

## Discussion

In this study, we have explored the effectiveness of UV light to inactivate the SARS-CoV-2 virion particles present in aerosols. UV radiation is long known to be sterilizing agent for microbes in the laboratories, hospitals, and food industry and is regularly used to sterilize various materials. Unlike bacterial, parasitic, or fungal pathogens, viruses has small genetic material with a simpler physical structure and thus are more susceptible for UV-induced destruction(5). UV-C is considered to be the most effective band of the UV spectrum and efficiently catalyzes the formation of photoproducts in DNA(23). Thymidine dimers irreversibly interrupt essential process of replication, transcription, and translation culminating into the inability of virus particle to establish infection. This study demonstrate that UV-C rapidly inactivates the SARS-CoV-2 virions present in the air.

SARS-CoV-2 is highly infectious virus which is evident from rapid spread of this virus from close or continuous contact with the infected person or from common touch surfaces. Our study using the air sampler, however, revealed that virus in the aerosols is quite sensitive to physical collision of the aerosols onto the solid surface and desiccation. Both these factors could be affecting the membrane integrity or Spike protein structure which has likely accounted for the virus inactivation. The results suggest that strong air current might help virus inactivation in the air and UV-C would further reduce the viral load to non-infectious baseline.

In 2020, when world witnessed huge COVID-19 infections and death toll, there were no vaccines and effective anti-viral drugs available. Upon the onset of vaccines in 2021, we are poised to have better control over the pandemic and are in far better condition to prevent the new infections or disease severity. However, new studies reveal that the vaccine-mediated immunity wanes over time and increases the chance of re-infection(24, 25). Moreover, the elderly and immunocompromised people are more susceptible to re-infections even after vaccination. Getting back to the normal life is important for the sustainability and will require effective method to inactivate the SARS-CoV-2 from our surrounding to prevent the chances of infection. As SARS-CoV-2 is air-borne and highly infectious rapid and effective air sterilization could substantially contribute to reduce viral load in air. In addition, the pathogens are not likely to acquire any resistance to this sort of mechanical inactivation. Our study reveals that UV-C radiation effectively inactivates SARS-CoV-2 in highly concentrated virus-loaded aerosols in the air. The exposure to the UVC radiation equivalent to 0.28 mJ/cm^2^ is enough to reduce >99% viruses and therefore can be used to device the strategies of air-purification.

In conclusion, the air-disinfection technology based on UV is a promising alternative to implement disinfection protocols in crowded places. The workplaces such as large offices, hospitals, etc are potential settings that are vulnerable for rapid transmission. The UV systems are generally used in combination with HEPA filters which surely enhances the pathogen filtering capacity but also can increase the cost and maintenance of the systems. The viral pathogens especially respiratory viruses could be inactivated by simple UVGI systems. The facile design of our systems presented in this study and the methodologies are accurate for sizing UV systems.

## Author Contributions

HG, SD, SP designed and fabricated the UV in-duct systems. KGT, AK, RPR designed experiments related to testing of in-duct systems to inactivate SARS-CoV-2 in aerosols. RPR and BT performed experiments related to SARS-CoV-2 live virus. KB and PBP did whole-genome sequencing of SARS-CoV-2 isolate. KK, Aku (Ashok Kumar), and NBB performed mathematical modelling using experimental data. HG and RR wrote the original draft of the paper with the comments from AK, KK, Aku.

## Acknowledgement

We thank Dr. Shekhar Mande, Dr Ram Vishwakarma, Dr Rakesh Mishra, Dr Anjan Ray, Dr Anantha Ramakrishna, Dr Sanjeev Khosla, Dr. N. Gopalakrishnan and other members of the CSIR Strategy Group (CSG) for COVID for their guidance and advice in conceiving this project. We are grateful to Dr. Hari Om Yadav for coordinating the project on air disinfection systems. We thank Arushi Goel and Nishant Raj Kapoor for technical assistance. This work was funded by grant from Council of Scientific and Industrial Research (CSIR) and Science and Engineering Research Board (IPA/2020/000168 to RPR). RPR is the recipient of DBT-Ramalingaswami fellowship. This is manuscript 035/2022 from CSIR-Institute of Microbial technology.

